# Familial thyroid follicular cell carcinomas in a large number of Dutch German longhaired pointers

**DOI:** 10.1101/2021.03.12.434920

**Authors:** Yun Yu, Adriana Krupa, Rebekah I. Keesler, Guy C. M. Grinwis, Mariska de Ruijsscher, Johan de Vos, Martien A. M. Groenen, Richard P. M. A. Crooijmans

## Abstract

Thyroid carcinomas originating from follicular cells of the thyroid gland occur in both humans and dogs and they have highly similar histomorphologic patterns. In dogs, thyroid carcinomas have not been extensively investigated, especially concerning the familial origin of thyroid carcinomas. Here we report familial thyroid follicular cell carcinomas confirmed by histology in 54 Dutch origin German longhaired pointers. From the pedigree, 45 of 54 histopathologically confirmed cases are closely related to a pair of first-half cousins in the past, indicating a familial disease. In addition, genetics contributed more to the thyroid follicular cell carcinoma than other factors by an estimated heritability of 0.62 based on pedigree. The age of diagnosis ranged between 4.5 and 13.5 years, and 76% of cases were diagnosed before 10 years of age, implying an early onset of disease. We observed a significant higher pedigree-based inbreeding coefficient in the affected dogs (mean *F* 0.23) compared to unaffected dogs (mean *F* 0.14), suggesting the contribution of inbreeding to tumour development. The unique occurrence of familial thyroid follicular cell carcinoma in this dog population and the large number of affected dogs make this population an important model to identify the genetic basis of familial thyroid follicular cell carcinoma in this breed and may contribute to the research into pathogenesis, prevention and treatment in humans.

## Introduction

Many dog breeds are predisposed to a variety of specific cancers due to consanguinity and inbreeding.^1^ According to research in the UK, cancer is the major cause of death in dogs, accounting for 27% of all deaths. Skin and soft tissues were the most common sites for tumour development, followed by alimentary, mammary, urogenital, lymphoid, endocrine and oropharyngeal.^2^ Within the tumours in the endocrine organs, thyroid carcinoma (TC) is the most common type, which represents 1.2% to 3.8% of all canine tumours and accounts for 90% of thyroid tumours.^3–5^ Thyroid carcinoma can originate from either follicular cells (follicular cell carcinoma, FCC) or parafollicular cells (C-cell carcinoma). Within FCC, four main histological subtypes of differentiated thyroid carcinomas (dTC) are described: follicular thyroid carcinoma (FTC), compact thyroid carcinoma (CTC), follicular-compact thyroid carcinoma (FCTC), and papillary thyroid carcinoma (PTC) with FTC and CTC the most frequent.^4,6^ Furthermore, poorly differentiated and undifferentiated carcinomas, and thyroid carcinosarcomas (TCs) are also recognized.^6^ In humans, TC is the ninth most common type of cancer and accounts for approximate 3.1% of all cancers.^7^ The histologic growth patterns in humans are largely similar to those in dogs. Additionally, TC shows no sex preference in dogs, although in humans, females have a 3-fold higher risk than males.^5,8^ The prevalence of TC in older dogs (between 10 and 15 years old) is significantly higher compared to earlier onset.^5^

Thyroid tumours can be of familial or spontaneous origin. In humans, the majority of TCs are sporadic and approximately 5-15% of them are considered to be of familial origin.^9,10^ Due to the relatively low prevalence of familial TCs, the genetic causes are less investigated than sporadic types, thus are still poorly understood.^11^ To the authors’ knowledge, in dogs, there has only been one pedigree of apparent familial medullary TC reported.^12^ Investigations and reports of familial thyroid tumours in dogs have been limited.

Over a period of more than 21 years, a relatively large number of TCs were diagnosed in the German longhaired pointers born in the Netherlands (Dutch GLPs). In this retrospective study, we review clinical and histopathological assessments of the GLPs with thyroid tumours, and present genetic assessment including the inbreeding and heritability estimation based on pedigree.

## Materials and Methods

### Study population

Medical records of the clinics belonging to Dutch and Belgian collaborating veterinary cancer centres and the database of two Dutch veterinary diagnostic pathology laboratories were searched for client-owned GLPs diagnosed with thyroid tumours between 1996 and 2017. Additionally, the owners of GLPs registered in the database of the Dutch GLP association were contacted to identify any dogs with a history of thyroid tumour. Once the dog was diagnosed with a thyroid tumour, the primary or referring veterinarian was contacted to obtain relevant information. If more than one dog was affected in the litter, the owners of the remaining littermates as well as dogs related to each of the parents were identified and contacted. Pedigree records were provided by GLP “Langhaar” association (www.germanlonghair.com) in order to perform a pedigree analysis.

Only GLPs with histopathologically confirmed follicular cell thyroid carcinoma were included as cases in this study and used in genetic analysis. Surgical removal of the affected thyroid glands, when feasible, was centralized in one clinic (AH Z-Vl).

Cases were excluded if owners rejected to participate in the research. All the data and samples in this research were permitted to be used for scientific purpose and in publication.

### Clinical data

The following information was retrieved from the medical records, if available: signalment, physical examination findings including tumour size (longest diameter), location and mobility (determined by palpation), clinical signs, time to presentation and date of diagnosis.

Whenever performed, the results of additional diagnostic tests, including blood tests and imaging tests, were recorded. Blood tests included complete blood cell count, serum biochemistry profiles, basal circulating total thyroxine (TT4) and thyroid stimulating hormone (TSH) concentrations. Staging was performed using diagnostic imaging (thoracic radiographs, cervical ultrasonography, computed tomography [CT]). If available, the presence of ectopic thyroid tumour was recorded.

### Histopathology analysis

For histopathological evaluation, tissues harvested during surgery or necropsy were fixed in 10% neutral buffered formalin. Representative sections were routinely embedded in paraffin and sectioned at 4 μm and stained with haematoxylin and eosin (H&E) and examined via immunohistochemistry for thyroglobulin and calcitonin expression.

Thyroglobulin and calcitonin immunohistochemistry (IHC). Four-micron tissue sections of the formalin-fixed and paraffin-embedded tumour tissue were dried overnight, deparaffinized and rehydrated with xylene (2×5min) and 100% alcohol (2×3min). Endogenous peroxidase activity was blocked with 1% H2O2 in methanol for 30 minutes. After rinsing in 1% Tween20 in PBS, the slides were treated with 1:10 normal goat serum in PBS for 15 minutes and incubated for 60 minutes with 1:200 diluted Rabbit anti-human thyroglobulin (Dako, Denmark) or 1:400 diluted Rabbit anti-human calcitonin (Dako, Denmark) at room temperature. After rinsing in PBS/Tween, the slides were then incubated with Goat anti-rabbit/biotin (Vector Labs) secondary antibody (dilution 1:250 in PBS) for 30 minutes at room temperature. The slides were rinsed with 1% Tween20 in PBS, incubated with ABC/PO complex (Vector Labs) for 30 minutes and rinsed with PBS. Lastly the slides were incubated with DAB solution for 25 minutes and counter stained with haematoxylin for 30-60 seconds at room temperature. For negative controls the primary antibody was omitted.

Tissues were evaluated by two veterinary pathologists and classified according to the World Health Organization (WHO) classification of tumours of the endocrine system scheme.^6^ If a tumour had multiple growth patterns, then classification was based on the most predominant pattern. If capsular penetration of the neoplasm was unclear, additional H&E sections were cut for additional evaluation.

### Genetic analysis

To assess the genetic relationship between dogs collected, the family tree of all unaffected and affected (suspected and histopathologically diagnosed) dogs were constructed using Kinship2 package in R.^13^ Pedigree based inbreeding coefficients (*F*) of all the dogs were estimated using the CFC program^14^ based on the whole pedigree of GLPs. To evaluate the contribution of inbreeding to the incidence of the thyroid tumours in the population, the rank sum test of *F* between affected dogs histopathologically confirmed and unaffected dogs born before 2007 was done in R using Wilcoxon test. We excluded 86 unaffected dogs born after 2007 because many of these dogs are closely related to the affected dogs, and they could be highly susceptible to FCC. Although they are unaffected at the time of analysis, they could become affected later in their lives, thus biasing the result.

### Heritability estimation

Heritability was estimated using ASReml 4.1 based on the pedigree relationship between the unaffected dogs and cases histopathologically confirmed.^15^ Unaffected dogs born after 2007 were also excluded from the estimation. The model used is as follows.

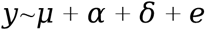

Where y is the phenotype, which is a binary trait, affected status coded as 1 and unaffected status coded as 0. α is the fixed effect of gender, female or male. δ is the random animal effect. e is the random residual.

Heritability calculation equation is:

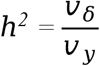

Where *ν_δ_* is the variance of the random animal effect, *ν_y_* is the variance of FCC phenotype.

## Results

In total, 264 GLPs born between 1991 and 2017 were identified (supplementary Table S1). One hundred eighty dogs were unaffected and had no signs of thyroid tumour at the time of entering the study, data analysis or during follow up (1996-2019). Twenty-nine dogs were suspected of thyroid neoplasia based on typical clinical signs like the presence of cervical mass, but no further diagnostics have been performed. These dogs were suspected cases in this study. Fifty-four dogs met the inclusion criteria of real cases given the histopathological diagnosis of FCC. One dog was additionally diagnosed with thyroid adenoma. Among the 54 cases, 34 (63%) were male (4 castrated, 30 intact) and 20 (37%) were female (7 spayed, 13 intact). The median age was of 8 years (range, 4.5-13.5 years). Forty-one dogs (76%) developed thyroid tumour before reaching the age of 10 years.

### Clinical complaints

Forty-four of 54 dogs (81%) had information regarding clinical complaints related to thyroid tumour recorded. Duration from the onset of clinical signs to the presentation ranged from 61 to 732 days. Detection of palpable thyroid mass without any other concurrent signs was reported in the majority of dogs (37). Seven dogs (13%) demonstrated additional clinical signs that included: intermittent cough (3 dogs), alopecia (3 dogs), polyuria (2 dogs) polydipsia (2 dogs), weight loss (1 dog), and lethargy (1 dog).

One dog was asymptomatic with the diagnosis of the first thyroid tumour but developed clinical signs at the time of contralateral tumour. In contrast, another dog presented with complaints related to the first thyroid tumour, while no clinical signs were recorded at diagnosis of the second tumour.

### Tumour details

Bilateral tumours were identified in 35 dogs, and unilateral tumours in 19 dogs. Eleven tumours were left-sided, 6 right-sided, and for 2 the site of involvement was not mentioned. Three dogs were suspected of having ectopic tumours: two in the cranial mediastinum, one at the base of the heart.

Of the 23 tumours for which information regarding the palpable mobility of the mass was available, 13 were described as moveable, whereas 10 were described as fixed. Mobility of the remaining tumours on palpation was not specified in the medical record.

Information regarding tumour size was most available in the form of the maximum dimension. Estimated tumour size based on physical examination was available for 33 dogs. Median maximal tumour diameter was 5 cm (range 2 to12 cm).

### Diagnostic findings

Forty-nine dogs (91%) had information regarding diagnostic imaging, including CT of the cervical region and thorax (13 dogs), cervical ultrasonography (3 dogs), thoracic radiographs (22 dogs) and abdominal ultrasonography (4 dogs).

Based on diagnostic imaging, 4 dogs had involvement of the regional lymph nodes: 2 dogs ipsilateral retropharyngeal lymph node, 1 dog ipsilateral mandibular and retropharyngeal lymph node and 1 dog ipsilateral cervical superficial lymph node. Histopathology confirmed metastatic disease in 3 dogs. One dog underwent post-mortem examination, but the suspected lymph node was not evaluated.

Distant metastases were suspected in only one dog (pulmonary nodules) however further diagnostics were not performed to confirm this.

### Clinical pathology

On presentation, TT4 (total T4) was measured in 30 dogs and TSH in 11 dogs. Four dogs with elevated TT4 and decreased TSH showed clinical signs compatible with hyperthyroidism. Seventeen dogs had TT4 within normal limits, while in 9 dogs it was below the lower end of the reference interval. Four dogs had elevated TSH while their TT4 was also increased (3 dogs) or within the reference ranges (1 dog). Three dogs had unremarkable TSH and TT4.

Other clinical pathological abnormalities were sporadic and mild, including anaemia (3 dogs), leukocytosis (1 dog), hypocalcaemia (1 dog), alkaline phosphatase elevation (1 dog), and hypercholesterolemia (4 dogs).

#### Histopathology

Thyroid follicular cell carcinomas were diagnosed in 54 dogs. Bilateral neoplasms were diagnosed in 29 dogs. The majority of the 83 carcinomas showed a follicular growth pattern (n=37; fig 1A), whereas compact (solid) (n=15; fig 1B), follicular-compact (n=16) and papillary (n=9; fig 1C) growth patterns were seen in the other carcinomas. In 3 dogs a carcinosarcoma, characterized by osteosarcoma and carcinoma (fig 1D), was diagnosed. In 2 dogs a carcinoma not otherwise specified (NOS) was diagnosed. In 1 dog diagnosed in 1996, which was the first case we found, the diagnosis was only thyroid tumour with signs of malignancy. In 4 carcinomas, well-differentiated bone tissue was seen (metaplastic bone formation). An ectopic compact follicular cell carcinoma was found at the heart- base during necropsy in one dog that also had follicular-compact type carcinoma in both thyroid glands.

**Fig 1.**
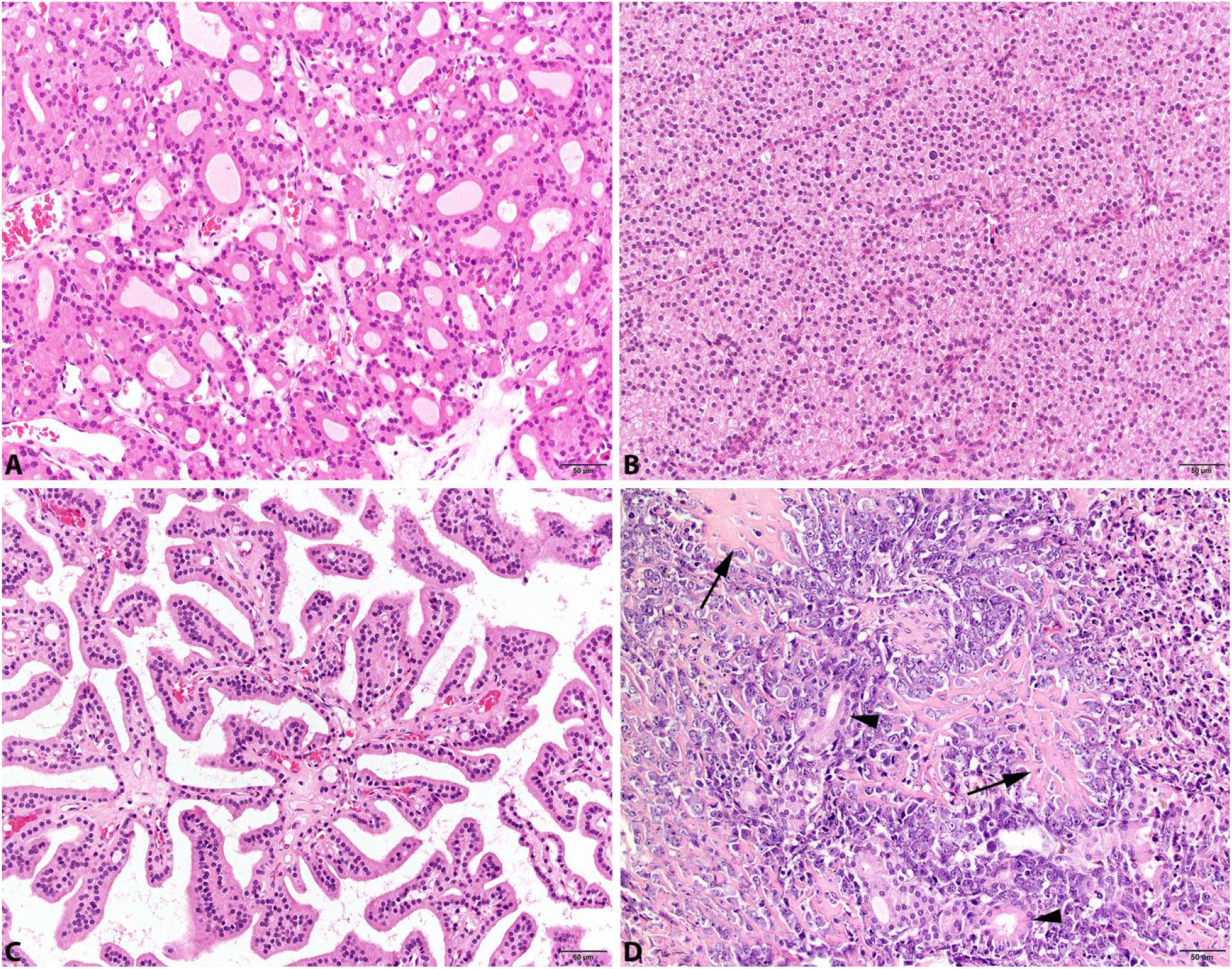
Histological pictures of different histological types of thyroid follicle cell carcinomas in German longhaired pointers. Follicular (A), compact (B) and papillary (C) growth pattern of neoplastic cells. D shows a carcinosarcoma with osteoid (arrows) producing mesenchymal neoplastic cells and scattered neoplastic follicular structures (arrowheads). H&E.

Immunohistochemistry was performed on the neoplasms of 40 dogs. The neoplastic cells were vaguely to markedly positive for thyroglobulin in all tumours. The strongest immunoreactivity was typically noted in the colloid with lower staining intensity in the neoplastic cells. All neoplasms were negative for calcitonin.

#### Heritability

For the heritability estimation besides the 54 histologically confirmed cases, 94 unaffected GLP dogs born before 2007 were incorporated in the analysis. Heritability of the FCC in these dogs was estimated to be 0.62 (±0.19).

#### Inbreeding

The complete GLP pedigree registered worldwide used for inbreeding estimation included 58,634 GLPs. The 17,786 Dutch GLPs have higher inbreeding coefficient (average *F*=0.19) compared to GLPs born in other countries (average *F*=0.10) with a *p*-value of 2.2e-16 (fig 2A). Based on this complete GLP pedigree, the inbreeding coefficients of 52 of 54 histologically confirmed affected dogs was 0.23 where in the unaffected dogs born before 2007 it was 0.14. Affected dogs are more inbred than unaffected dogs (*p*-value 2.473e-08) (fig 2B).

**Fig 2.**
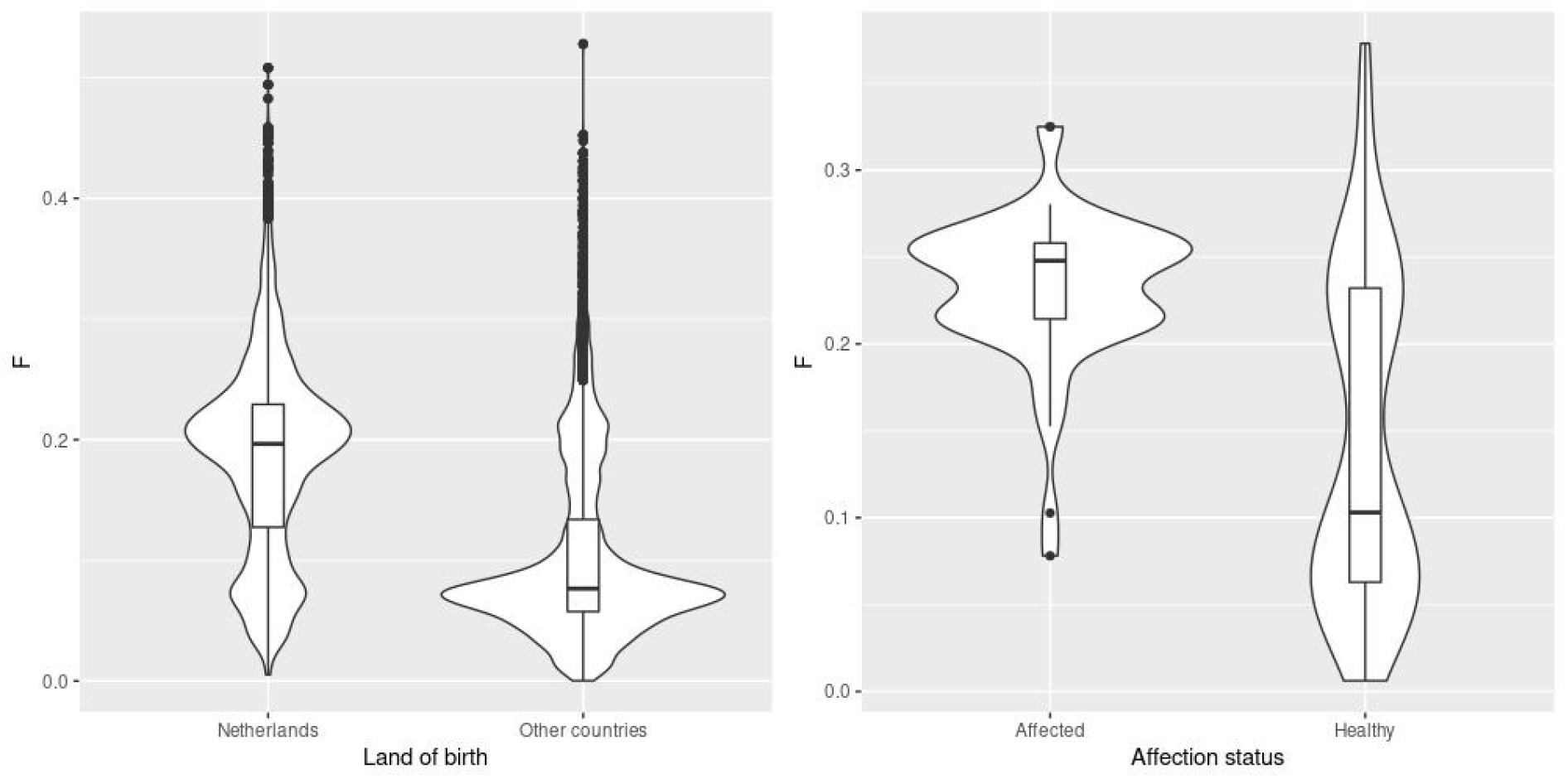
(A), Inbreeding of dogs born in the Netherlands and other countries. (B), Inbreeding of histopathologically confirmed affected GLPs and unaffected GLPs born before 2007.

#### Relationship between affected dogs

According to the pedigree of all collected GLPs (supplementary fig 1), most affected dogs are very closely related. Meanwhile, a strikingly high incidence of FCC was observed in some families due to intensive use of a few prominent dogs. Forty-five affected dogs are related to one pair of first-half cousins GLP52 and GLP905 (fig 3) with a relationship coefficient of 0.21. Twenty-four affected dogs are the first generation of offspring of GLP52 (47 dogs in total) with an incidence of the TC of 51%. Twenty-two affected dogs are the first generation of offspring of GLP905 (140 dogs in total) where incidence reaches 16%. Moreover, five affected dogs descended from siblings of GLP52, and four affected dogs are descendants of siblings of GLP905. GLP52 is a suspected case and has one suspected affected full-sibling GLP53. GLP905 has an unknown case status due to inaccessibility, but has one suspected affected full-sibling, GLP13, and one affected half-sibling, GLP47, with histological diagnosis.

**Fig 3.**
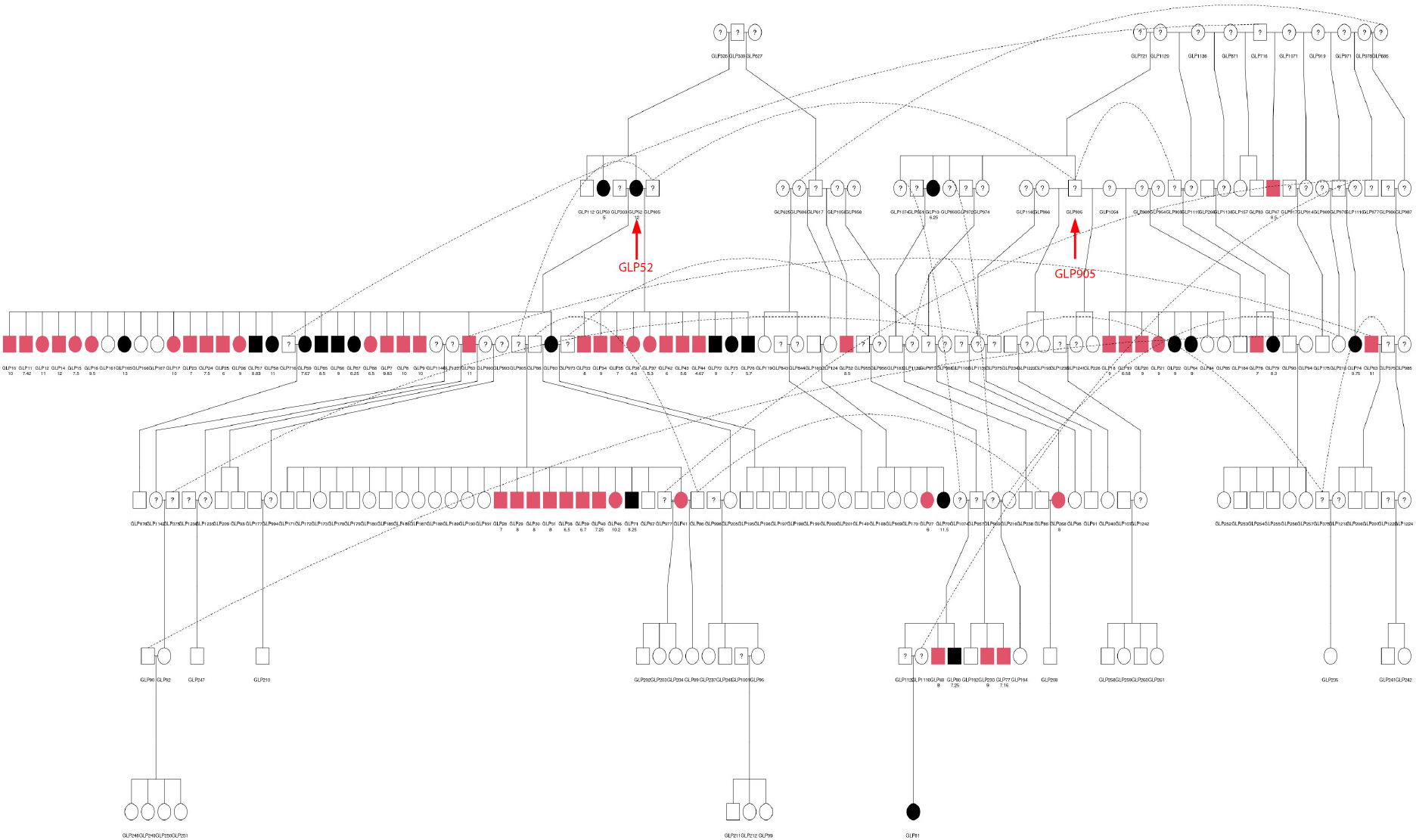
Pedigree of dogs related to GLP52, GLP905. Forty-five histopathologically confirmed affected dogs are closely related to these two dogs. Affected dogs with histological diagnosis are highlighted in red, and suspected affected dogs (without histopathology diagnosis) are in black, whereas unaffected dogs remain white. A question mark represents the dogs with unknown status.

## Discussion

In humans, familial TC is diagnosed when two or more first-degree relatives are affected.^10^ Here, we showed that the incidence of FCC is strikingly high in some families of Dutch GLPs, like in the pedigrees of GLP52 and GLP905. These two dogs have a most recent common ancestor, GLP306 (supplementary fig 3), born in 1989, with the *F* of 20.57%. Furthermore, 78 probably affected GLPs (26 suspected and 52 histopathologically confirmed FCC cases) can be traced back to a common cross of 6 generations prior to GLP52, the cross between GLP319 and GLP296 (supplementary fig 2). With such close relationships between the majority of the affected dogs, the FCC in these dogs is considered to be a familial disease.

In this study, besides the 54 histopathologically confirmed cases, twenty-nine dogs were suspected to be affected by thyroid tumour based on clinical findings (e.g., presence of a mass lesion at the location of the thyroid gland), but because no histological assessment was performed, these suspicions could not be confirmed. Interestingly, these suspected cases are very closely related to the most affected GLPs with diagnosis (supplementary fig 5). Among them, twenty-two are closely related to the two prominent spreaders of the disease, GLP52 and GLP905 (fig 3), as either the siblings or direct descendants. These suspected dogs are very likely affected by the same familial FCC.

Familial cancers usually occur at a relatively young age. Thyroid carcinoma normally occurs at the median age of 9-10 years in dogs and its occurrence increases with age.^3^ In a previous study, approximately 57% of FTC in dogs occurred between 10 and 15 years,^5^ while in our cohort of Dutch GLPs, the FCC showed early onset with 76% of cases occurring before 10 years of age. However, some cases can have very late onset, as there are 10 dogs with an age at diagnosis of > 10 years, which could represent spontaneous cases within our cohort.

In humans, familial thyroid follicular cell cancer has an autosomal dominant inheritance pattern with incomplete penetrance.^16,17^ However, FCC in these Dutch GLPs is likely a recessive trait, according to the occurrence of FCC in the family of GLP160 and GLP124 (fig 4). GLP124 is the offspring of a half-sibling of GLP52, and GLP124 and GLP160 have a common ancestor with GLP52 and GLP905, a male dog born in 1971. Both GLP124 and GLP160 were unaffected, while one of their 5 offspring was confirmed to be affected and one was a suspected case. This confirms the recessive behaviour of the trait, although considering the possible incomplete penetrance of this disease, there is a small chance that unaffected parents could be carriers of a dominant causal gene but do not show the phenotype. Therefore, to determine completely whether the TC in this study is recessive or dominant, further genetic analysis is needed.

**Fig 4.**
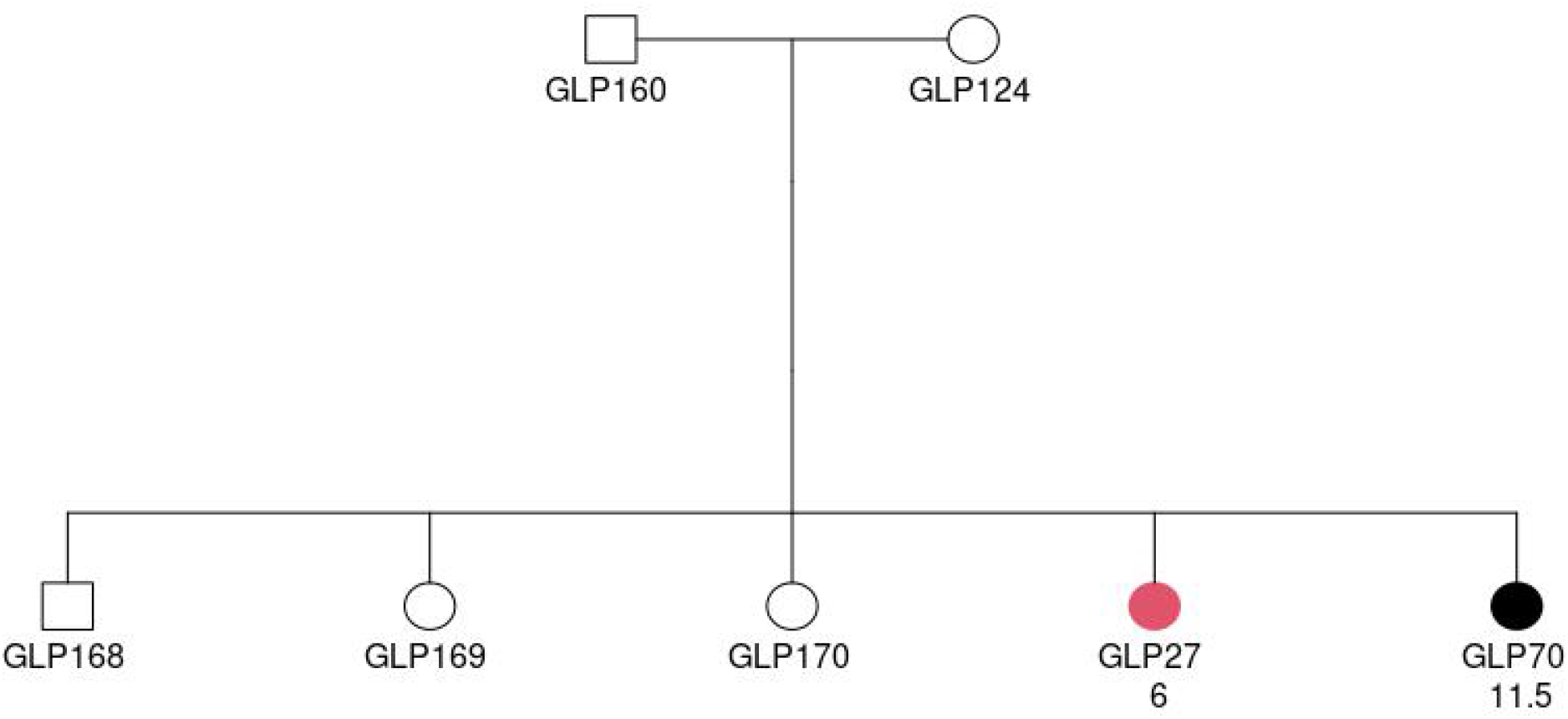
Affected status of a cross between unaffected individuals GLP160 and GLP124. Square denotes male and circle represents female. The individual in red was confirmed to be affected by histopathology. Black colour indicates a suspected case based on clinical signs. The 2 rows of texts below the circles or squares represent the ID and diagnosis age (in years), respectively.

One reason why thyroid tumours occurred in so many GLPs is that the age of detection is higher than the typical breeding age. Before any signs of the thyroid tumours were noticed, the dogs produced offspring, like GLP52. GLP52 was a dog affected at 12 years of age, but that had already been crossed with GLP905 and GLP333 and produced 37 affected offspring, many years before the first case was diagnosed in the offspring generation. In addition, intensive use of few dogs in the breeding programs also contributed to the high incidence of TC in these dogs. In total, GLP52 and GLP905 have 602 and 512 descendants, respectively, demonstrating how a few dogs were used intensively. These dogs and their future offspring all have high susceptibility to TC because of the consanguinity.

Inbreeding is an important tool used in dog breeding programs to fix desirable traits in a population. However, harmful side effects, such as inbreeding depression, could decrease animal performance and result in a high risk of propagation of recessive diseases or defects,^18,19^ as demonstrated in this study. Inbreeding contributed to the high incidence of FCC in this dog population, because we found a significant higher *F* in the affected GLPs compared to the unaffected GLPs (fig 2B). Moreover, the 2 prominent spreaders, GLP52 and GLP905, are highly inbred, with inbreeding coefficients of 0.21 and 0.24, respectively. Both parents of GLP52 are from inbred crosses between half-siblings. We also see other extreme inbreeding examples which produced affected dogs. For instance, GLP905 was crossed with its half-sibling GLP1119 and produced 2 affected dogs (1 confirmed, 1 suspected).

Cancer incidence is complex and is determined by a combination of many factors, including genetic make-up, the environment and the lifestyle of the carrier, with genetics playing a large role. In humans, thyroid carcinoma has the strongest genetic component among all the cancers, with genetic contribution exceeding other factors.^20^ In these GLPs with TC, genetic factors may contribute more than environmental factors as well, with a heritability estimated to be 0.62.

The genetic basis of familial thyroid cancer is poorly defined in humans, as only 5% of familial FCC cases have well-defined germline mutations.^11^ Research of thyroid carcinoma in dogs can contribute to the knowledge of corresponding thyroid carcinoma in humans. Dogs have been proposed as an ideal model for human cancer research, because many cancers have strong similarity in histological appearances, genetic causes, biological behaviours and response to conventional therapy.^21^ Additionally, dogs share their environments with human pet owners, thus are partly exposed to similar risk factors, which can be exploited for epidemiological studies of cancers common in humans and dogs.^22^ The affected GLPs we reported here can serve as a model to identify the genetic basis of FCC. We have a uniquely large number of affected dogs from one breed, and they are inbred (average *F* 0.23) and very likely share common genetic mutations that are associated with carcinogenesis. The large sample size gives more possibility and power to further define the underlying mutation(s) of this disease by genetic and genomic techniques, like e.g., whole genome association analyses.

## Acknowledgement

We thank the members of the SKK working group of the Langhaar association for providing access to the dog breeders and owners including financial support for the research. We especially thank Annie van der Sluis for her help in the pedigree data collection. Yun’s PhD study was supported by scholarship #201806760044 from China Scholarship Council (CSC).

## Author contributions

Ada Krupa and Johan de Vos performed clinical diagnosis and analysis. Rebekah I. Keesler and Guy C.M. Grinwis performed histopathological analysis. Mariska de Ruijsscher and Johan de Vos collected the data. Richard R. P. M. Crooijmans and Yun Yu designed the study. Yun Yu wrote the manuscript with input from all authors. Johan de Vos and Richard Crooijmans set up the early ideas of this research.

## References

1. Schiffman J, Breen M. Comparative oncology: What dogs and other species can teach us about humans with cancer. Philosophical transactions of the Royal Society of London Series B, Biological sciences. 2015;370.

2. Dobson JM, Samuel S, Milstein H, Rogers K, Wood JL. Canine neoplasia in the UK: estimates of incidence rates from a population of insured dogs. J Small Anim Pract. 2002;43(6):240–246.

3. Barber LG. Thyroid Tumors in Dogs and Cats. Veterinary Clinics of North America: Small Animal Practice. 2007;37(4):755–773.

4. Liptak JM. Canine Thyroid Carcinoma. Clinical Techniques in Small Animal Practice. 2007;22(2):75–81.

5. Wucherer KL, Wilke V. Thyroid Cancer in Dogs: An Update Based on 638 Cases (1995— 2005). Journal of the American Animal Hospital Association. 2010;46(4):249–254.

6. Kiupel M, Capen C, Miller M, Smedley R. Tumors of the thyroid. Histological Classification of Tumors of the Endocrine System of Domestic Animals 2nd Series (Schulman, FY ed), Armed Forces Institute of Pathology, Washington, DC. 2008:25–39.

7. Rahib L, Smith BD, Aizenberg R, Rosenzweig AB, Fleshman JM, Matrisian LM. Projecting cancer incidence and deaths to 2030: the unexpected burden of thyroid, liver, and pancreas cancers in the United States. Cancer Res. 2014;74(11):2913–2921.

8. Bray F, Ferlay J, Soerjomataram I, Siegel RL, Torre LA, Jemal A. Global cancer statistics 2018: GLOBOCAN estimates of incidence and mortality worldwide for 36 cancers in 185 countries. CA: A Cancer Journal for Clinicians. 0(0).

9. S. AA, Kannan MA, Ebtesam Q, Hindi A-H. HABP2 Gene Mutations Do Not Cause Familial or Sporadic Non-Medullary Thyroid Cancer in a Highly Inbred Middle Eastern Population. Thyroid. 2016;26(5):667–671.

10. Nosé V. Familial thyroid cancer: a review. Modern Pathology. 2011;24:S19.

11. Ngeow J, Eng C. HABP2 in Familial Nonmedullary Thyroid Cancer: Will the Real Mutation Please Stand Up? JNCI: Journal of the National Cancer Institute. 2016;108(6).

12. Lee J-J, Larsson C, Lui W-O, Höög A, Von Euler H. A dog pedigree with familial medullary thyroid cancer. International journal of oncology. 2006;29(5): 1173–1182.

13. Sinnwell JP, Therneau TM, Schaid DJ. The kinship2 R package for pedigree data. Human heredity. 2014;78(2):91–93.

14. Sargolzaei M, Iwaisaki H, Colleau JJ. CFC: A tool for monitoring genetic diversity. Proceedings of the 8th World Congress on Genetics Applied to Livestock Production. 2006:27–28.

15. Gilmour AR, Gogel, B. J., Cullis, B. R., Welham, S. J. and Thompson, R. ASReml User Guide Release 4.1 Structural Specification, H. VSN International Ltd. 2015.

16. Kebebew E. Hereditary Non-medullary Thyroid Cancer. World Journal of Surgery. 2008;32(5):678–682.

17. Saporito D, Brock P, Hampel H, et al. Penetrance of a rare familial mutation predisposing to papillary thyroid cancer. Familial Cancer. 2018;17(3):431–434.

18. Doekes HP, Veerkamp RF, Bijma P, de Jong G, Hiemstra SJ, Windig JJ. Inbreeding depression due to recent and ancient inbreeding in Dutch Holstein—Friesian dairy cattle. Genetics Selection Evolution. 2019;51(1):54.

19. Ujvari B, Klaassen M, Raven N, et al. Genetic diversity, inbreeding and cancer. Proceedings of the Royal Society B: Biological Sciences. 2018;285(1875):20172589.

20. Czene K, Lichtenstein P, Hemminki K. Environmental and heritable causes of cancer among 9.6 million individuals in the Swedish Family-Cancer Database. Int J Cancer. 2002;99(2):260–266.

21. Khanna C, Lindblad-Toh K, Vail D, et al. The dog as a cancer model. Nature Biotechnology. 2006;24(9):1065–1066.

22. Rowell JL, McCarthy DO, Alvarez CE. Dog models of naturally occurring cancer. Trends Mol Med. 2011;17(7):380–388.

