## Supplemental Figure S1 - S5 for "Familial thyroid follicular cell carcinomas in a large number of Dutch German longhaired pointers"

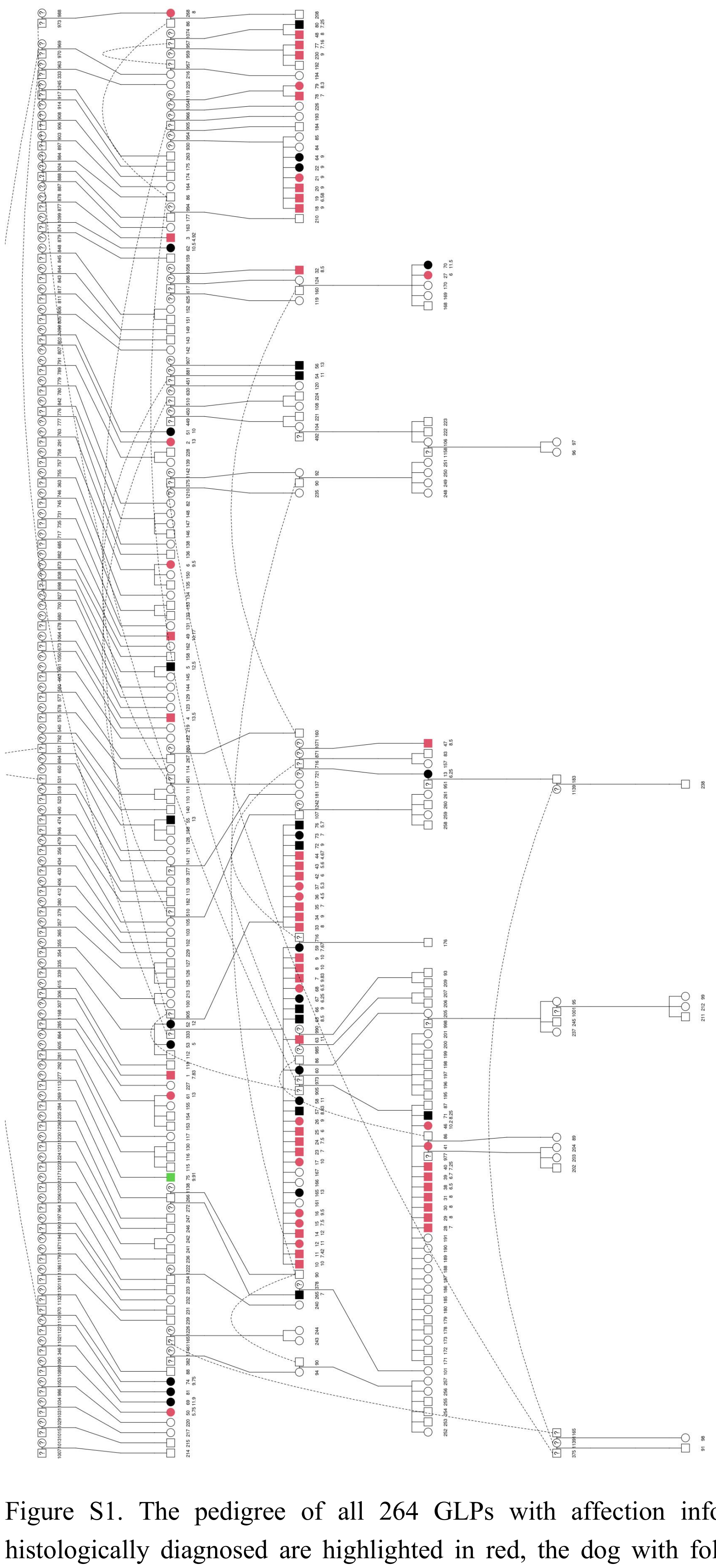

Figure S1. The pedigree of all 264 GLPs with affection information. Dogs with FCC histologically diagnosed are highlighted in red, the dog with follicular thyroid adenoma is highlighted in green, and suspected affected dogs are in black, whereas unaffected dogs remain white. A question mark represents the dogs with unknown status.

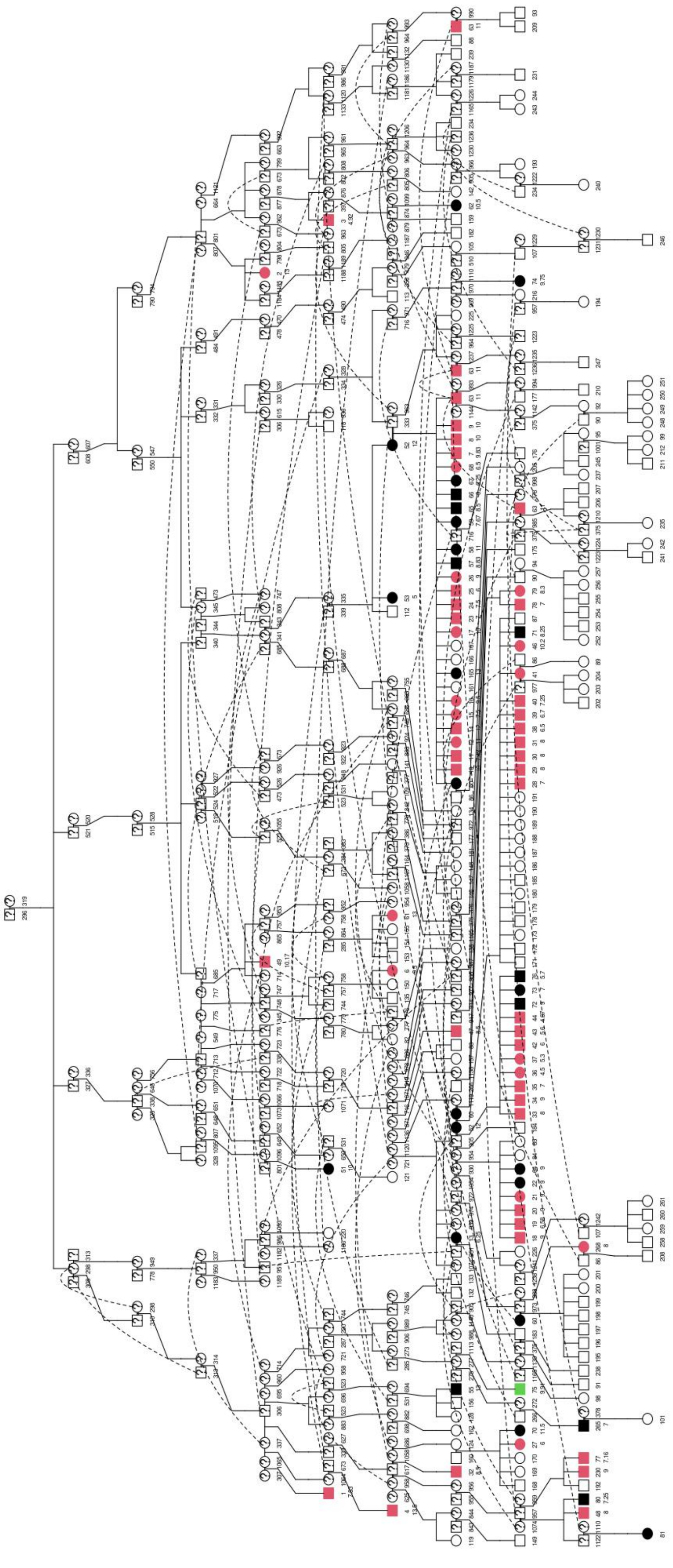

Figure S2. Pedigree of GLPs that could be traced back to the cross between GLP296 and GLP319. Dogs with FCC histologically diagnosed are highlighted in red, the dog with follicular thyroid adenoma is highlighted in green, and suspected affected dogs are in black, whereas unaffected dogs remain white. A question mark represents the dogs with unknown status.

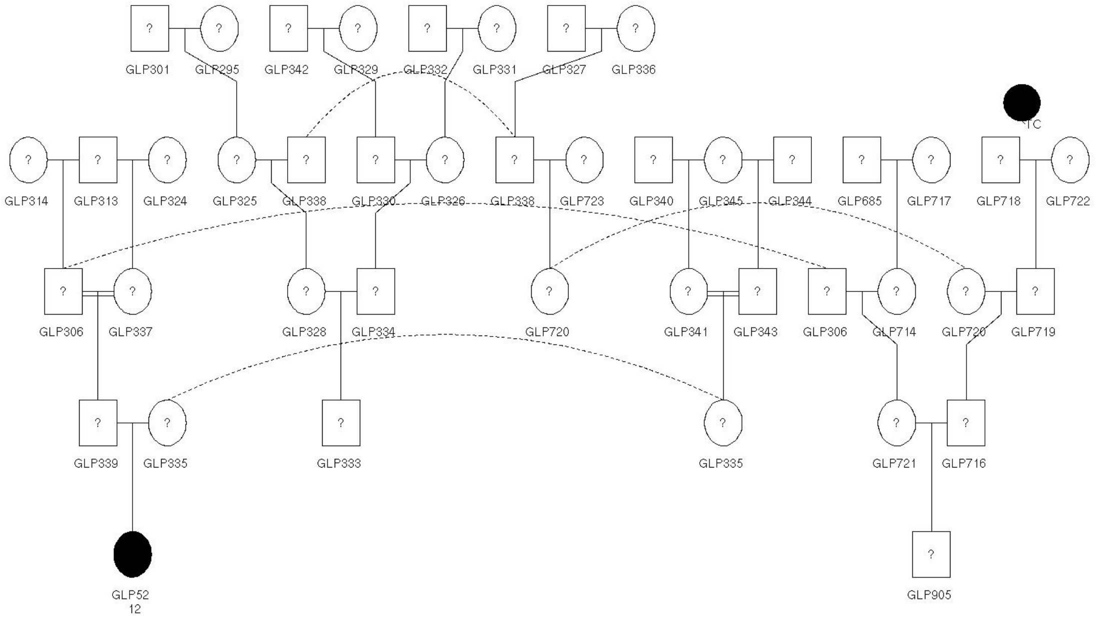


Figure S3. Ancestry family tree of GLP52 and GLP905. These two dogs are half-first cousins with a common grandfather GLP306. GLP52 was a suspected case.

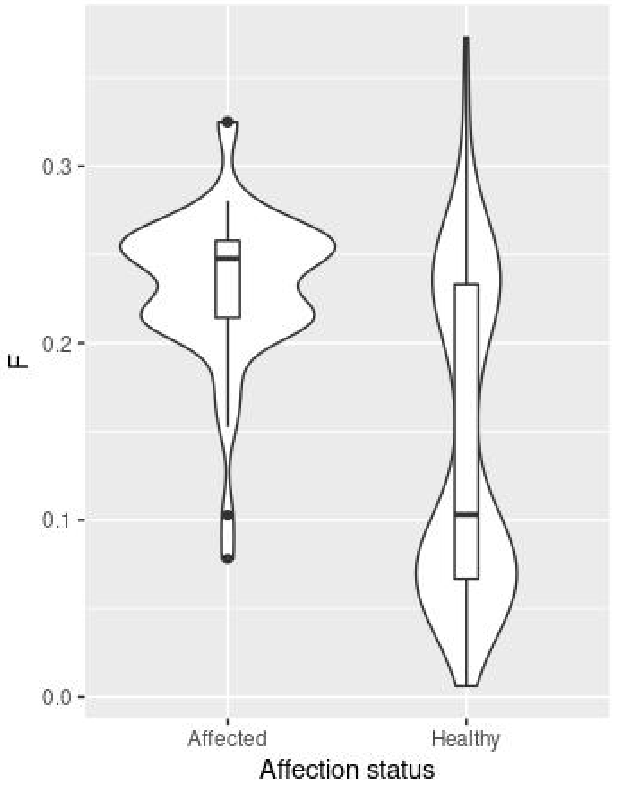


Figure S4. *F* of affected and unaffected GLPs including the dogs born after 2007 in our dataset (54 cases and 177 controls)(Wilcoxon test, *p*-value=4.317e-10). Affected dogs are more inbred than unaffected dogs.

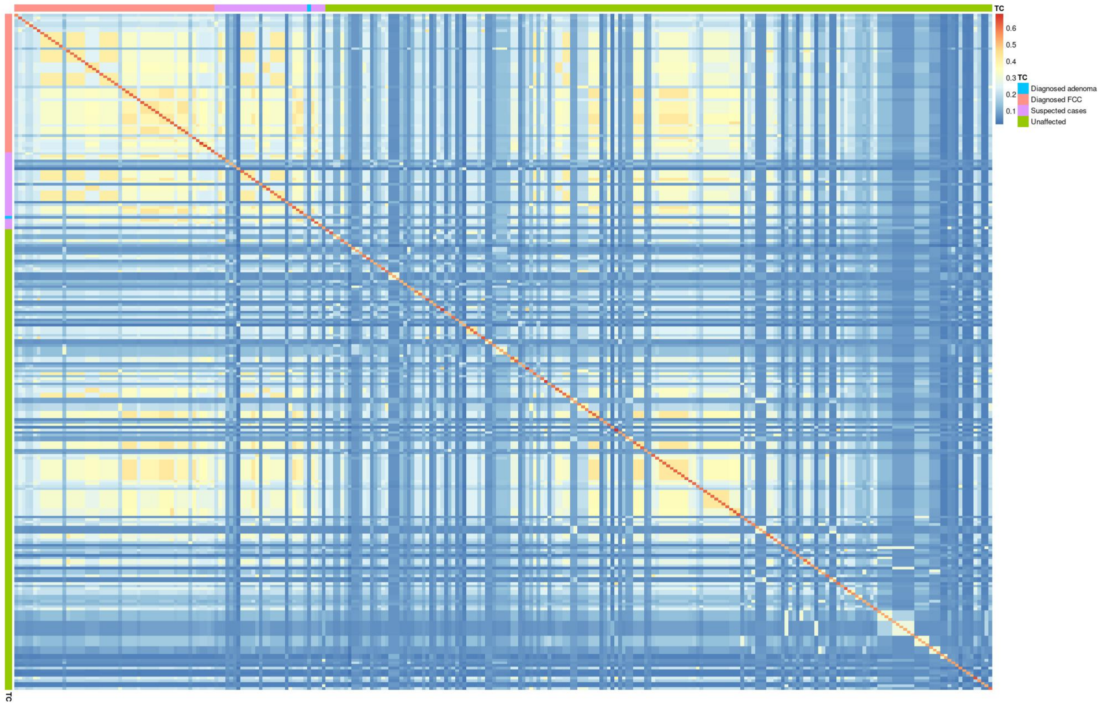


Figure S5. Kinship matrix between 54 FCC cases, 1 dog with adenoma, 29 suspected cases, and 180 unaffected dogs. Kinship matrix was estimated using kinship2 package in R. The 54 FCC cases are closed related to each other. Meanwhile, most suspected cases are closely related to the histologically diagnosed FCC cases.
